# Elemental Composition and Degradation Rate Impact the Biocompatibility of Copper Chalcogenide Nanocrystals

**DOI:** 10.64898/2025.12.17.695045

**Authors:** Xingjian Zhong, G. Perry Katsarakes, Savani Nagarkar, Allison M. Dennis

**Author notes:** Corresponding Author: Allison M. Dennis.

## Abstract

Copper chalcogenide nanocrystals (NCs) are promising candidates for biophotonic applications due to their tunable optical properties. Concrete methods to examine the relationship between their degradation and toxicity are necessary to enable development of nanoconstructs with reduced toxicity. This study compares the degradation and acute cytotoxicity of three compositions of micelle-coated copper chalcogenide NCs: the fluorescent semiconductor copper indium sulfide (CuInS_2_), and the plasmonic semiconductors copper sulfide (Cu_2-x_S) and chalcopyrite copper iron sulfide (CuFeS_2_). We developed a quantitative degradation assay to assess ion release from these ultra-small nanocrystals, revealing that while all three particles biodegrade, CuInS_2_ and CuFeS_2_ undergo rapid degradation in artificial lysosomal fluid, leading to a burst release of indium and iron ions. In cellular toxicity assays, CuInS_2_ exhibited significantly higher acute cytotoxicity than Cu_2-x_S and CuFeS_2_, primarily due to indium-induced necrosis. To mitigate this toxicity, an alternative surface-binding polymer coating was introduced, effectively reducing both the degradation rate and cytotoxicity of CuInS_2_. These findings highlight the influence of both nanocrystal composition and coating chemistry in moderating the acute cytotoxity of degradable nanocrystals, demonstrating that tuning of composition and degradation rate can be used to moderate nanoparticle toxicity.

## 1. Introduction

Copper chalcogenide-based semiconductor nanocrystals (NCs) are powerful photoactive agents for biomedical applications due to their tunable optical properties. Nanocrystals with composition Cu_x_M_y_S_z_, where M represents substituted metal elements, function as either direct bandgap photoluminescent semiconductor NCs or plasmonic semiconductor NCs.^1–3^ Photoluminescent indium-doped copper chalcogenide NCs, i.e., copper indium sulfide (CuInS_2_) quantum dots (QDs),^4–7^ have a size-, crystal structure-, and stoichiometry-dependent band gap that facilitates tunable emission throughout the visible wavelength range and into the near-infrared, including the first tissue optical window for bioimaging applications.^4,8^ Plasmonic copper chalcogenide NCs include vacancy-doped copper sulfide (Cu_2-x_S) and iron-doped copper iron sulfide (Cu_x_Fe_y_S) NCs, whereby elemental dopants and vacancies result in excess holes or electrons.^9–13^ These off-stoichiometry carrier concentrations lead to optical absorbance features from the localized surface plasmon resonance between incident light and the NCs.^2,14^ By tuning the cation composition, thus controlling carrier density, one can shift the plasmonic absorbance peak without changing the particle physical structure for absorbance-based biophotonic applications such as photoacoustic imaging, photothermal therapy, and photodynamic therapy.^15– 20^

These copper chalcogenide NCs are attractive for biomedical applications due to their heavy metal-free composition, and their generally assumed biodegradability. For example, studies have claimed that Cu_2-x_S NCs are superior to biostable gold nanoparticles for photoacoustic and photothermal applications, citing non-toxic biodegradation and excretion as a mitigation to long-term biocompatibility concerns of gold nanoparticles.^19,21,22^ In a more nuanced example, CuInS_2_ quantum dots (QDs) with a ZnS shell have been favorably discussed as an alternative to lead- and/or cadmium-containing QD compositions for fluorescence imaging applications, and their biocompatibility has been noted as providing a potential path for translational applications.^6,23–25^ However, in a previous study, we demonstrated that while zinc sulfide (ZnS) shelled CuInS_2_ quantum dots are biostable and well tolerated, they also accumulate in the liver of mice precluding straightforward translation.^26^ In contrast, bare CuInS_2_ quantum dots degrade in biological conditions and are largely excreted within 28 days, but exhibit toxicity *in vitro* and *in vivo*.^26^ Notably, the acute toxicity following CuInS_2_ injection in mice was severe, but resolved over time with *in vivo* markers of toxicity (e.g., organ index, liver enzyme activity) returning to normal levels faster than the copper and indium was excreted. These observations led us to hypothesize that both the degradation products *and* the degradation rate (which determines local ion concentration) are key determinants of biocompatibility or toxicity for biodegrading, i.e., excretable, inorganic nanomaterials.

In the previous study, toxicity could not be explicitly correlated with a specific compositional element (Cu or In) as the concentration of copper and indium in the CuInS_2_ particles was proportional. To tease out the impact of Cu, In, and Fe on copper chalcogenide biocompatibility, we designed a comparative study of the degradation and acute cytotoxicity of similarly sized Cu_2-x_S, CuInS_2_, and CuFeS_2_ nanocrystals, each encapsulated in an FDA-approved PEGylated lipid micelle to provide water solubility. To assess which elements were being released in biological environments, we developed an accessible method to quantify degradation and link the rate and composition of the released cations to cytotoxicity results. Our compositional comparison shows that indium is a major contributor to toxicity, and the rapid release of indium from CuInS_2_ NCs in lysosomal environments likely cause significant cytotoxicity via cell necrosis. By changing the organic coating on the surface of the CuInS_2_ QDs, however, we demonstrate that wrapping the CuInS_2_ NCs in a surface-chelating polymer appears to slow the degradation of CuInS_2_ compared to lipid-PEG micelle encapsulation in simulated body fluid. This polymer-wrapped CuInS_2_ exhibited significantly reduced acute cytotoxicity compared to micelle-encapsulated CuInS_2_, suggesting that moderating degradation rate can reduce burst release of toxic ions. These results collectively demonstrate a strategy to quantify the degradation of inorganic nanoparticles and a proof-of-concept that coating chemistry can modulate both degradation kinetics and acute cytotoxicity. By demonstrating that both inorganic composition and organic coating influence degradation and acute toxicity, this work points to design strategies that may help address both degradation-related acute toxicity and long-term bioaccumulation concerns in nanomaterials. These insights should motivate further work toward excretable, biocompatible nanomaterials that leverage both compositional and coating strategies.

## 2. Results and Discussion

### 2.1 Three compositions of micelle-encapsulated copper chalcogenide nanocrystals

Three compositions of copper chalcogenide NCs were colloidally synthesized via hot injection reactions in an air-free environment and characterized using absorbance spectroscopy, transmission electron microscopy (TEM), and X-ray diffraction (XRD) (Fig. 1). The absorbance spectra are normalized to peak values across the wavelength range displayed for visualization and comparison. CuInS_2_ NCs are direct bandgap QDs and exhibit a broad absorbance spectrum that increases in intensity at higher energy wavelengths. Because CuInS_2_ nanocrystals are non-emissive due to surface oxidation without a ZnS shell, we rely on absorbance measurements for optical measurements. The CuInS_2_ NCs have an average size of 3.9 ± 0.6 nm measured with TEM images (Fig. S1). CuInS_2_ shares a similar chalcopyrite crystal structure with the size-matched CuFeS_2_ NCs (3.6 ± 0.7 nm; Fig. S1). Unlike CuInS_2_, CuFeS_2_ NCs exhibit a broad localized surface plasmon resonance (LSPR) feature in the visible wavelength range with a peak around 500 nm due to excess carriers from iron doping.^27,28^ Cu_2-x_S NCs were synthesized with a 2:1 Cu:S stoichiometry; upon exposure to air, unavoidable oxidation leads to the loss of some Cu ions, resulting in vacancies and excess hole-doping, which generates an LSPR absorbance feature in the NIR region (Fig. S2),^10,15^ seen as a rising tail towards 800 nm in Fig. 1. The Cu_2-x_S NCs are slightly larger than the CuInS_2_ and CuFeS_2_ NCs at 10.7 ± 2.4 nm (Fig. S1). While all of these small particles yield characteristically broad XRD peaks, comparison to reference peaks shows that the crystal structure of Cu_2-x_S aligns with a mixed phase of hexagonal djurleite (Cu_1.94_S) and cubic digenite (Cu_1.8_S), and align with the expected peak positions for chalcopyrite CuInS_2_ and CuFeS_2_.

**Figure 1.**
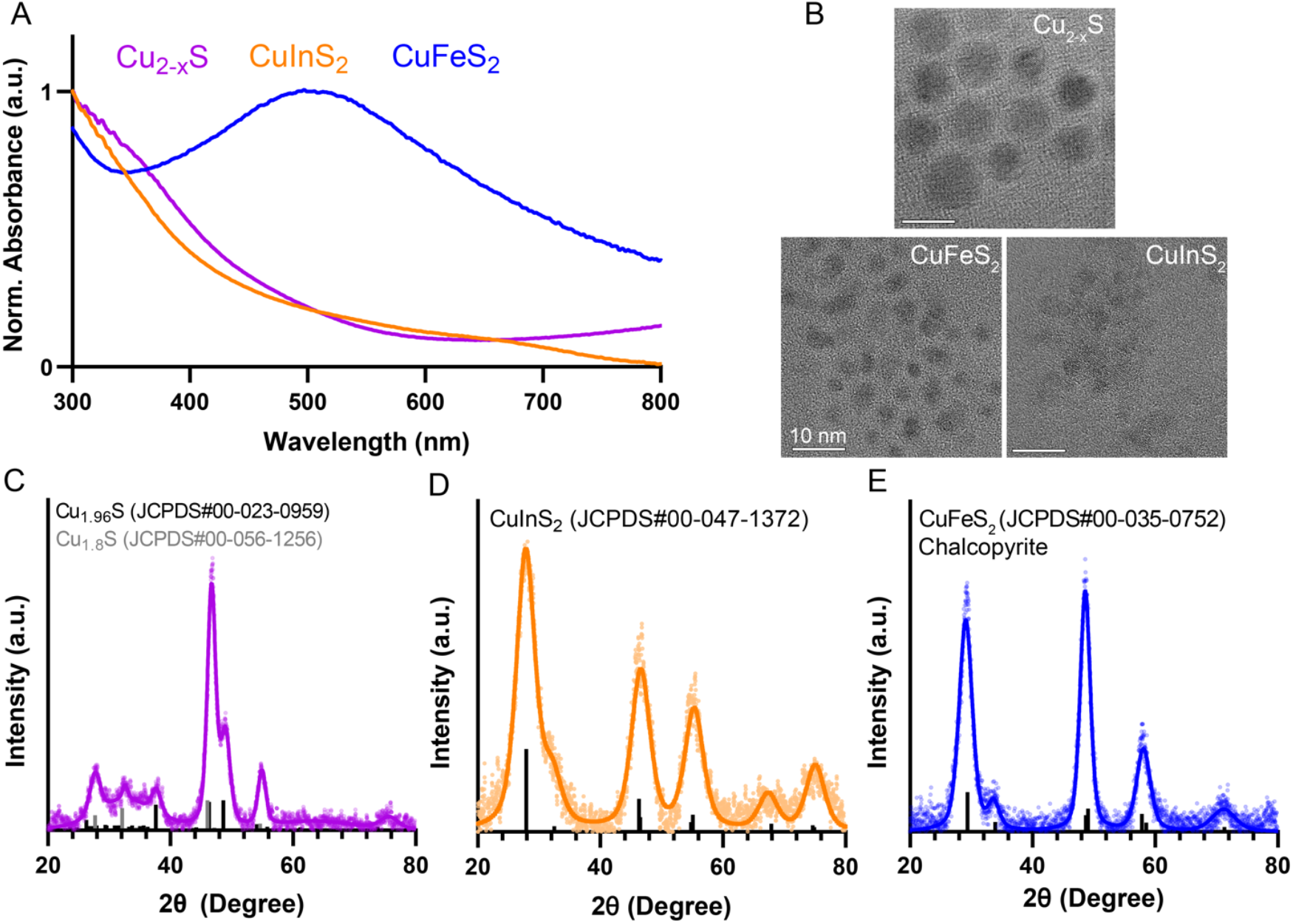
Characterization of copper chalcogenide nanocrystals. A) Absorbance spectra of CuInS_2_, CuFeS_2_ and Cu_2-x_S normalized to the maximum intensity in the range 300-800 nm. B) Representative TEM images of the three particles. (scale bar = 10 nm) C – E). XRD spectra of the particles fitted with reference peaks from International Centre for Diffraction Data (ICDD) database.

Each particle was synthesized in organic phase solutions with octadecene as the non-coordinating solvent and coordinating ligands oleylamine, trioctylphosphine, and/or oleic acid serving to stabilize the colloids in apolar liquids. For water solubility and utilization in biological environments, the NCs were coated with the FDA-approved PEGylated lipid DSPE-PEG (i.e., 1,2-distearoyl-sn-glycero-3-phosphoethanolamine-N-[amino(polyethylene glycol)2000], DSPE-PEG_2k_), encapsulating each particle in a lipid-PEG micelle through hydrophobic interactions between the hydrocarbons on the organic ligands on the surface of the NCs and the lipids. The micelle-encapsulated NCs were separated from free lipids, empty micelles, and aggregates using density gradient ultracentrifugation. All three NCs were encapsulated in the same biocompatible lipid-PEG coating to ensure consistency in both the NC-coating and coating-media interfaces for the degradation and toxicity comparisons. The physiochemical similarities between micelle coated NCs including size and surface charge allow for comparative studies focused on particle composition (Fig. S3).

### 2.2 Degradation analysis of the chalcogenides

Upon successful synthesis and characterization of the three copper chalcogenide compositions and their micelle encapsulation, we investigated their degradation behavior in biologically relevant environments. We initially assessed the degradation of the micelle-encapsulated particles with repeated absorbance spectroscopy measurements during incubation in simulated body fluid (SBF) and artificial lysosomal fluid (ALF) at 37°C (Fig. 2). SBF and ALF mimic key biological environments in their salt compositions and pH, while avoiding biomolecular confounders.^26,29,30^ Biological factors such as biomolecules and enzymes may alter degradation kinetics *in vivo*; however these defined buffer systems allow controlled comparison of degradation behavior across nanocrystal compositions. We observed degradation via loss in the high energy absorbance intensity for all of the NC compositions and a loss of signature absorbance features for CuFeS_2_ and CuInS_2_, indicated by a reduction in LSPR absorbance peak and the first excitonic feature near 700 nm, respectively. We also observed an elevated baseline absorbance for CuInS_2_ and CuFeS_2_ in SBF (Fig. 2A & B), likely caused by scattering from aggregated particles, which arises due to loss of colloidal stability, with or without degradation of the semiconductor nanoparticle. From the absorbance measurements, we infer that each of the tested copper chalcogenides degrades in biological conditions, but these results do not assess degradation quantitatively. Furthermore, given the distinct optical features of the various NC compositions, we were unable to identify a single optical parameter that would enable direct comparison between the materials. While these qualitative absorbance measurements confirmed degradation, we required a more precise method to quantify and compare compositional ion release dynamics across the different nanocrystal compositions.

**Figure 2.**
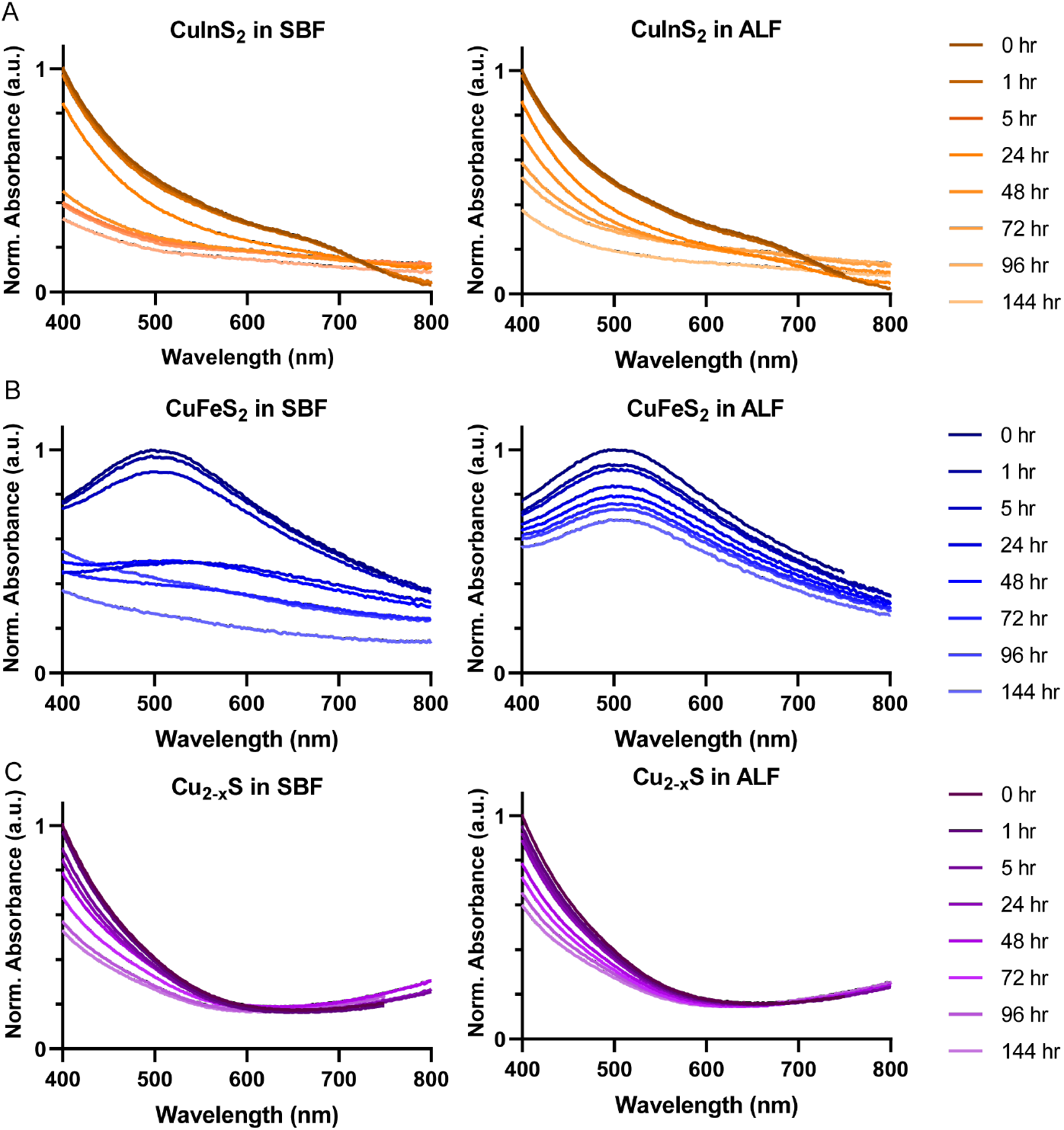
Qualitative degradation of micelle-encapsulated NCs observed via absorbance spectroscopy. (**A**) CuInS_2_, (**B**) CuFeS_2_, and (**C**) Cu_2-x_S incubated in simulated body fluid (SBF) and artificial lysosomal fluid (ALF). Absorbance spectra are normalized to the maximum initial absorbance intensity for each NC (i.e., t = 0 hr).

To quantify degradation, we devised a more direct assessment of ion release from these NCs by separating the particles and released ions, but we noticed that the limited methods to quantify degradation of ultra-small NCs were expensive, relatively inaccessible, and had not been applied to copper chalcogenides.^31,32^ For microparticles, separation is easily achieved via centrifugation using a bench-top centrifuge, however in the case of these small, colloidally stable NCs, ultracentrifugation with long run-time is necessary to physically separate the particles through centrifugation (e.g., >80 hr at >150,000 × *g* to form a hard pellet^33^). One could alternatively use centrifugal filters or dialysis membranes to physically separate intact particles from released ions, but this approach is frustrated by the high affinity between cations and cellulose membranes: our tests using metal salt solutions indicated that > 70% of Fe and In and ∼20% of Cu bound to cellulose dialysis membranes over a 4 hr incubation period (data not shown). Bespoke methods requiring rare or expensive instrumentation have been developed to study some materials. For example, iron oxide nanoparticle degradation was quantified using electron paramagnetic resonance spectroscopy to detect the ferromagnetic resonance correlated with intact particle mass.^31^ This method is not easily accessible nor applicable for the three copper chalcogenides evaluated in this study, so we found it necessary to develop a new assay for the quantitative analysis of NC degradation.

To separate intact particles from released ions, we precipitated the lipid-PEG coated NCs with ethanol, aggregating the particles, and pelleting them with bench-top centrifugation, leaving the soluble, released ions in the supernatant. The pellets and dried supernatants were digested with high purity nitric acid, diluted with ultrapure water, and the elemental compositions measured with MP-AES (Fig. 3A). The method was tested in pilot studies by comparing MP-AES results from fresh, intact particles and particles preemptively digested with nitric acid. For example, for CuFeS_2_ NCs, we recovered > 95% of cations from the intact particles in the pellet with an immeasurable amount in the supernatant, while > 95% of the cations from predigested CuFeS_2_ in acidic solution were recovered in the supernatant with trace amounts of copper or iron found in the pellet.

**Figure 3.**
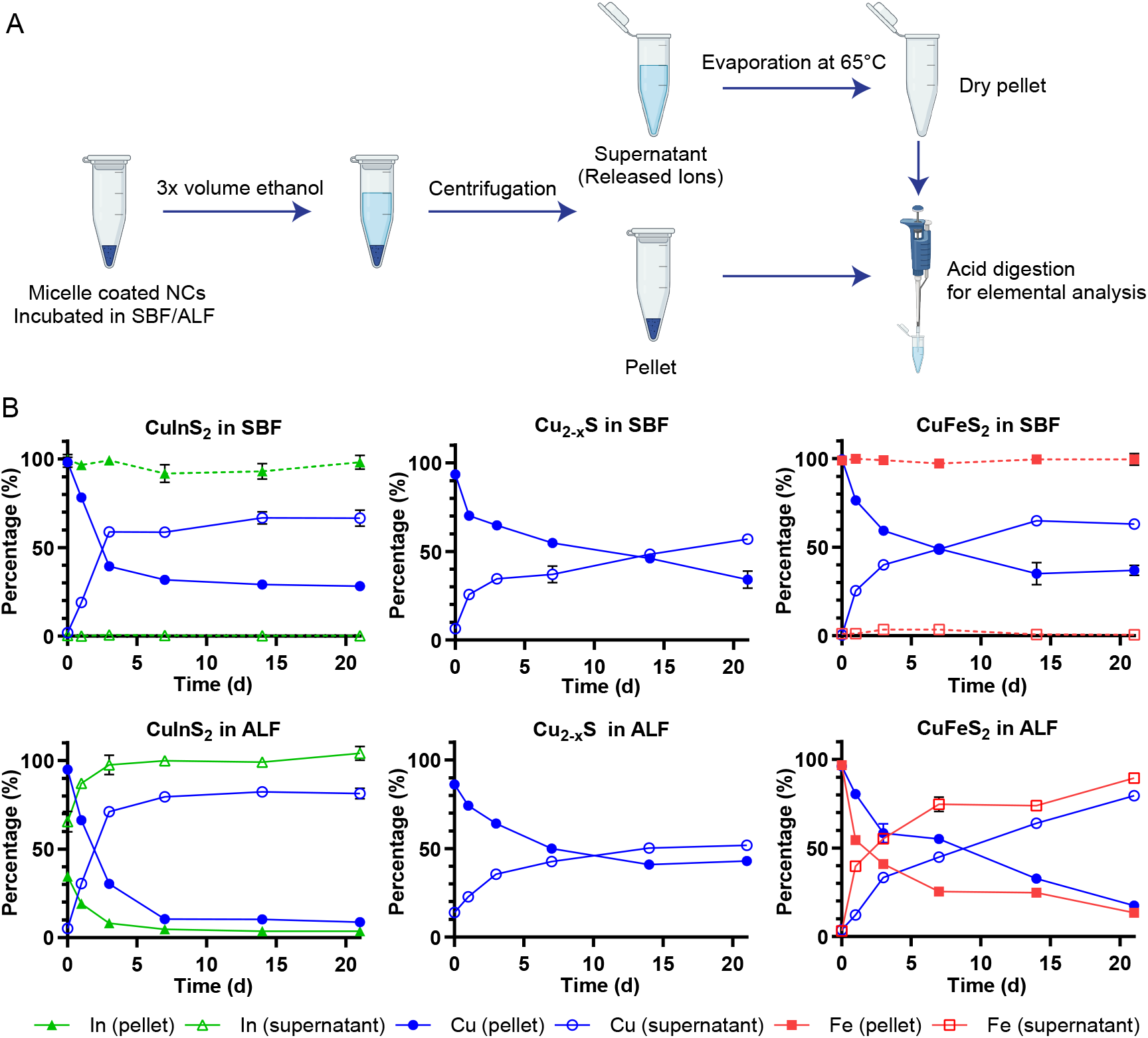
Quantitative degradation assay. (**A**) Scheme depicting the degradation experiments for micelle-encapsulated NCs. After NCs are incubated in the specified buffer, the soluble, released ions are separated from the persisting particle through ethanol precipitation and centrifugation. The separated samples are digested with nitric acid and analyzed for their elemental composition with microwave plasma atomic emission spectroscopy (MP-AES) (Schematic created with biorender.com). (**B**) Quantitative elemental analysis data normalized to total particle concentration for NCs incubated in ALF and SBF (n = 3). Ion species are represented in red rectangles, green triangles, and blue circles for Fe, Cu and In, respectively; solid and hollow shapes represent the pellet and supernatant, respectively. The earliest timepoint (t ≈ 10 min) represents the minimum time for sample preparation and measurement following dilution in SBF or ALF.

When comparing different compositions, the measurements from NCs incubated in ALF or SBF at different time points were normalized to the initial particle concentrations and plotted with respect to time in Figure 3B. For all experimental groups, we observe a monotonic increase in the release of Cu ions from the NCs over time, as shown through the increasing fraction of the Cu ions found in the supernatant of the separated particles. CuInS_2_ exhibited faster copper release in both ALF and SBF compared to the other compositions, with 59% and 71% of copper found in the supernatant for the CuInS_2_ samples at 3 d, respectively, compared 33-40% of copper in the supernatant for the CuFeS and Cu_2-x_S samples (Fig. S4). CuFeS copper release was comparable to CuInS_2_ at later timepoints, with both CuIn_2_S and CuFeS showing higher copper ion release in ALF compared to SBF at 21 d.

Although we can confirm the degradation of CuFeS_2_ and CuInS_2_ in SBF via absorbance measurements (Fig 2), the Fe and In from the SBF samples are consistently and completely located in the pellet in our elemental analysis. Buffer components such as phosphate can coordinate and facilitate release of Fe^3+^ and In^3+^ from the nanocrystals, but at neutral pH these ions form insoluble hydroxide complexes^34,35^ that precipitate with the particle aggregates. In contrast, the most striking results are seen in the elemental analysis of the In and Fe found in the supernatant following brief incubation in ALF. In the acidic, chelating environment of ALF, we see rapid Fe release and an almost immediate In release from the particles. In both cases, the Fe/In release outpaces the release of Cu into the supernatant. We attribute the fast Fe release to the chelation of Fe^3+^ by citric acid in the ALF, as described in other studies.^31,36^ We suspect the same chelation effect likewise drives the release of In^3+^, yielding >60% In^3+^ release in the sample taken within minutes of the NCs being dispersed in ALF (t ≈ 10 min ), >85 % release within 24 hr, and complete In^3+^ release by day 3 (Fig 3, S4).

### 2.3 Cytotoxicity of Micelle-encapsulated NCs

After establishing the distinctive degradation profiles of the three copper chalcogenide compositions, we proceeded to investigate how these degradation characteristics correlate with cellular toxicity. We first performed cell viability studies on the HepG_2_ cell line to compare the acute cytotoxicity between NCs. This liver cell line was chosen because a majority of nanoparticles in vascular circulation accumulate in the liver.^37^ Cell viability data are presented against total cation concentration (Cu+In, Cu+Fe, or Cu alone) to enable direct comparison of ionic toxicity across different species. Note that sulfur is not expected to impact cell viability and is challenging to quantify using standard elemental analysis. Following 24-hr incubation with micelle-coated NCs, CuInS_2_ exhibited the most significant cytotoxicity with a half-maximal inhibitory cation concentration (IC_50_) of 120 μg/mL, while Cu_2-x_S NCs were better tolerated with an IC_50_ of 230 μg/mL. CuFeS_2_ induced the least cell death, and cell viability was 65% after 24 hr incubation with our highest concentration of CuFeS_2_ at 1.2 mg/mL Cu+Fe (Fig. 4A). The cell viability results are consistent with our previous results for micelle-encapsulated CuInS_2_.^26^ When comparing CuInS_2_ and CuFeS_2_ with the same crystal structure, particle size, and micelle coating, we observe that the degradation profile is very similar, but the cytotoxicity is considerably higher for CuInS_2_ than CuFeS_2_.

**Figure 4.**
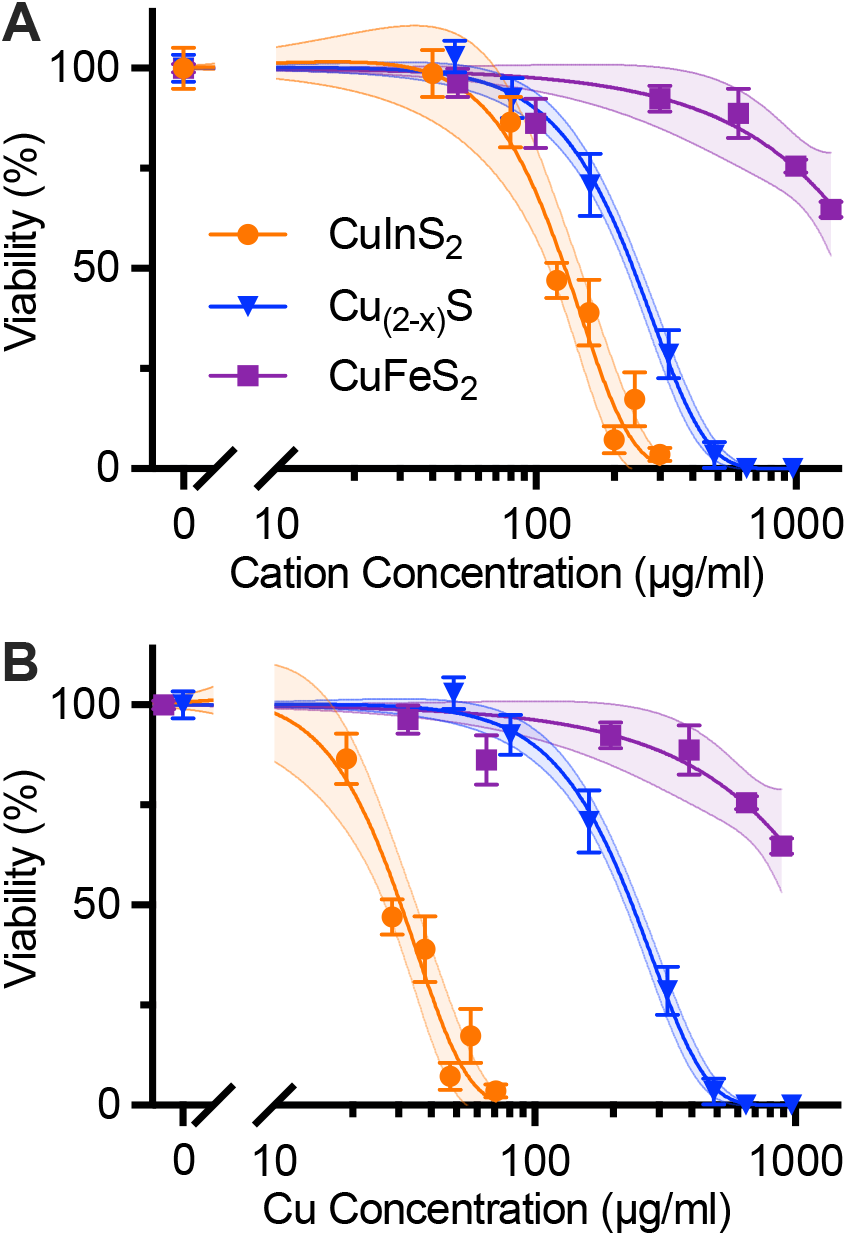
Cell viability of HepG_2_ liver cell line after 24-hr incubation with NCs. Data are normalized to negative control and fitted to a sigmoidal dose response curve. 95% confidence intervals of the fits are shown as the shaded areas around each curve. (**A**) Cell viability of HepG_2_ cells incubated for 24 hr with micelle-coated NCs and assayed with CellTiter-Glo. The fit curves yield IC50_cation_ values of 120 mg/mL and 230 µg/mL total cation concentration (i.e., Cu+In or Cu) for CuInS_2_ and Cu_2-x_S, respectively. (**B**) Data from A plotted against Cu concentration, yielding IC50_Cu_ values of 30 µg/mL and 220 µg/mL for CuInS_2_ and Cu_2-x_S, respectively. IC50s for CuFeS_2_ not reported, because cell viability was 65% at the highest tested doses of 1360 µg/mL and 890 µg/mL cation and copper concentrations, respectively.

When comparing CuInS_2_ to Cu_2-x_S, indium appears to have a larger influence on cell viability than Cu. This is further shown by re-plotting the cell viability data against Cu concentration for all three particle compositions (Fig. 4B). The gap widens between the response profiles of CuInS_2_ and Cu_2-x_S, indicating that indium is a more direct source of toxicity from CuInS_2_ than copper. While differences in surface reactivity between the various nanocrystal compositions could contribute to the observed toxicity variations, the stark contrast in toxicity between CuInS_2_ and CuFeS_2_—despite their similar structure, size, and degradation patterns—provides compelling evidence that indium plays the dominant role in the observed cytotoxicity.

While both In^3+^and Fe^3+^ both exhibit rapid release in the acidic and chelating conditions of ALF, only Fe is a bioessential element; both pH and chelation play a key role in iron homeostasis in cells and whole mammals.^38,39^ Biological systems have evolved sophisticated pathways to manage essential metals like iron, copper, and zinc through specialized transporters and storage mechanisms. These systems tightly regulate intracellular concentrations, preventing toxic accumulation.^40^ In contrast, non-essential metals like indium lack these regulatory mechanisms, leaving cells vulnerable to toxicity when exposed to sudden increases in ionic concentrations. This fundamental difference in biological handling likely explains why the released indium ions cause more severe cellular damage than iron ions, despite their similar charges and release kinetics. Likewise, the rapid release of indium ions in a biological milieu like ALF could explain the acute toxicity we previously observed for CuInS_2_ *in vivo*.^26^

### 2.4 Moderating CuInS_2_ degradation with a wrapped polymer surface coating

Since the rapid In^3+^ release from the micelle-encapsulated CuInS_2_ NCs may be directly driving the increased cytotoxicity of these particles, we hypothesize that slowing the degradation of the CuInS_2_ particles, and moreover the release of the In^3+^ ions, may moderate the cytotoxicity by reducing the instant dose of the offending ions. To slow the rate of degradation, we coated the CuInS_2_ particles with a surface binding polymer, which exhibits a stronger interaction between the coating and the NC compared to the micelle-encapsulated particles, and monitored their stability through longitudinal absorbance measurements. A histamine-modified poly(isobutylene-*alt*-maleic anhydride) (PIMA) coating was introduced as an alternative to the lipid-PEG coating for the CuInS_2_ NCs with the hypothesis that the polymer wrapping could stabilize the CuInS_2_ NC against degradation. The PIMA polymer binds directly to the CuInS_2_ surface through chelation of the surface metal ions with the imidazole rings on the histamines, replacing the native organic ligands such as oleylamine and trioctylphosphine.^41–44^ A quaternary amine functional group ((2-aminoethyl) trimethylammonium) was added to balance the charge of the polymer and ensure a zwitterionic surface coating,^41,42^ resulting in a similar surface charge to the lipid-PEG encapsulated CuInS_2_ (Fig. S3). Modified PIMA polymers have been used in a number of drug delivery research applications because they are versatile platforms for introducing functional groups,^42,45,46^ and related polymers such as poly(methyl vinyl ether-*alt*-maleic anhydride) (PVME-MA), often known as Gantrez, are recognized by the FDA for their safety in various applications.^47^ Some concerns have been raised that the high density of negatively charged carboxylic acid groups on native PIMA may induce cytotoxicity;^48,49^ however, the charge-balancing zwitterionic formulation used here may mitigate this concern, though systematic evaluation of coating-related toxicity will be important in future work. The PIMA-coated CuInS_2_ (CuInS_2_-PIMA) exhibits a comparable hydrodynamic diameter and ζ-potential as the micelle-encapsulated CuInS_2_ (CuInS_2_-micelle; Fig. S3), enabling direct comparison of how the surface coating may impact acute cytotoxicity.

When comparing the micelle-encapsulated and PIMA-coated CuInS_2_, the PIMA coating appears to protect the NCs from degradation in SBF. Longitudinal absorbance measurements show that CuInS_2_-PIMA largely maintained the high energy absorbance intensity from intact NCs over the course of the week in SBF, while a slightly elevated baseline indicates some colloidal aggregation during the experiment (Fig. 5A). In comparison, the CuInS_2_-micelle exhibited a 60% reduction in their high energy absorption intensity within 2 d in SBF (Fig. 2A and 5B). Both coatings provide colloidal stability in aqueous dispersions, but the direct surface chelation by PIMA appears to better protect against particle dissolution compared to the physically encapsulating lipid-PEG micelle. The low pH of ALF protonates the imidazole rings used for polymer chelation to the NC surface, destabilizing the PIMA coating and resulting in aggregation, which precluded degradation measurements in the artificial lysosomal environment. The CuInS_2_-PIMA NCs were dispersible in cell culture media, however, enabling comparison of the acute cytotoxicity of both coatings up to 250 μg/mL with HepG_2_ cells (Fig. 5C). PIMA-coated CuInS_2_ exhibited significantly reduced acute cytotoxicity compared to micelle-encapsulated CuInS_2_, suggesting that slowing nanocrystal degradation can moderate the burst release of indium ions and reduce cellular toxicity. These results provide proof-of-concept that coating chemistry can modulate acute toxicity and warranted further evaluation with CuInS_2_-PIMA particles in a follow up apoptosis/necrosis assay.

**Figure 5.**
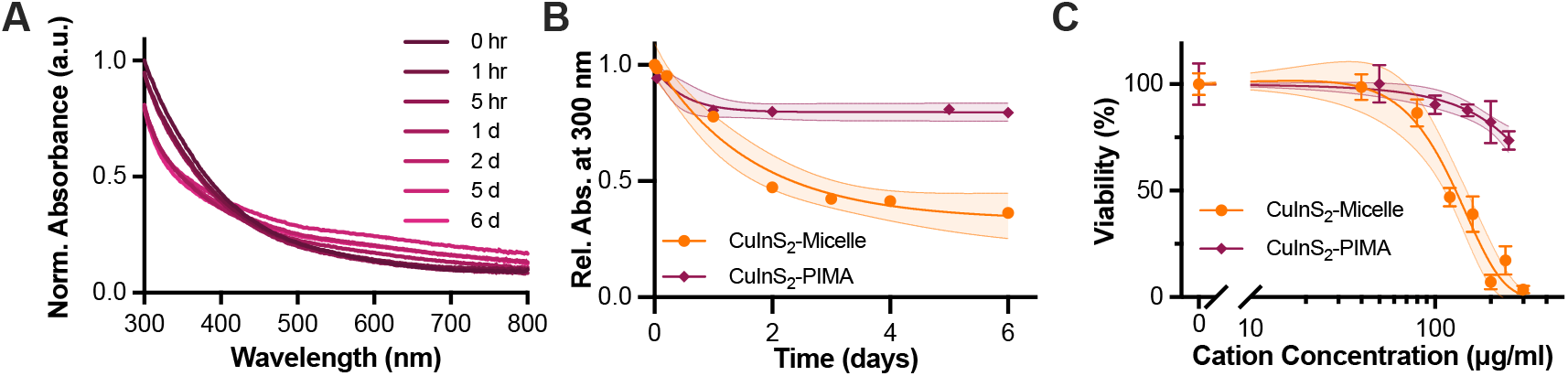
Comparing polymer-wrapped and micelle-encapsulated CuInS_2_. (**A**) Absorption of CuInS_2_-PIMA normalized to 300 nm at t_0_. (**B**) Comparison of high energy absorbance of PIMA-coated and micelle-encapsulated CuInS_2_ in SBF at 37 °C, indicative of NC stability and persistence, fit to a single exponential decay. (**C**) Comparison of cell viability of HepG_2_ liver cell line after 24-hr incubation with NCs. Data are normalized to negative control and fitted to a sigmoidal dose response curve. 95% confidence intervals of the fits are shown as the shaded areas around each curve.

### 2.5 Changes to composition or coating mitigate NC-induced necrosis

A multicolor fluorescence apoptosis/necrosis assay kit analyzed with flow cytometry was used to elucidate whether the NC-induced cytotoxicity was due to apoptosis or necrosis. HepG_2_ cells were incubated with 150 μg/mL cation concentration of micelle-encapsulated NCs and CuInS_2_-PIMA for 6 hr at 37°C. Following particle dosing and incubation, cells were detached and stained for 30 min according to the assay protocol. Stained cells were measured with flow cytometry, and the two channel results were presented with nuclear green DCS1 dye intensity representing membrane integrity, an indirect biomarker for necrosis, and the apopxin deep red indicator highlighting the membrane-bound early apoptosis biomarker (Fig. 6A). Cell culture media, ethanol, and the apoptosis-inducing drug staurosporine were used as control groups (Fig. S6). Since HepG_2_ are adherent cells, the assay results in measurable necrotic and apoptotic populations even in the negative control group, likely due to damage to cell membranes when detaching. From the flow cytometry results, we observed a substantial increase in the necrotic population from micelle-encapsulated CuInS_2_ relative to both the CuInS_2_-PIMA group and other NC compositions (Figure 5B). Cu_2-x_S also exhibited a slight increase in the necrotic population, while the CuFeS_2_ and CuInS_2_-PIMA groups were not statistically different from than the negative control. Among all apoptosis measurements, only the micelle-coated CuInS_2_ group showed a statistically significant increase compared to controls. However, most apoptotic cells in the micelle-coated CuInS_2_ group were found in the double-positive region, indicating they were simultaneously undergoing necrosis (Fig. S6). Overall, the HepG_2_ cell death correlates with NC-induced necrosis. By comparing the compositions as well as the different CuInS_2_ surface coatings, we link cell necrosis to the rapid degradation of CuInS_2_-micelle NCs and the burst release of indium ions, while replacing the indium with iron (CuFeS_2_) or slowing down the degradation (CuInS_2_-PIMA) significantly reduced the necrosis and hence the cytotoxicity (Fig. 7).

**Figure 6.**
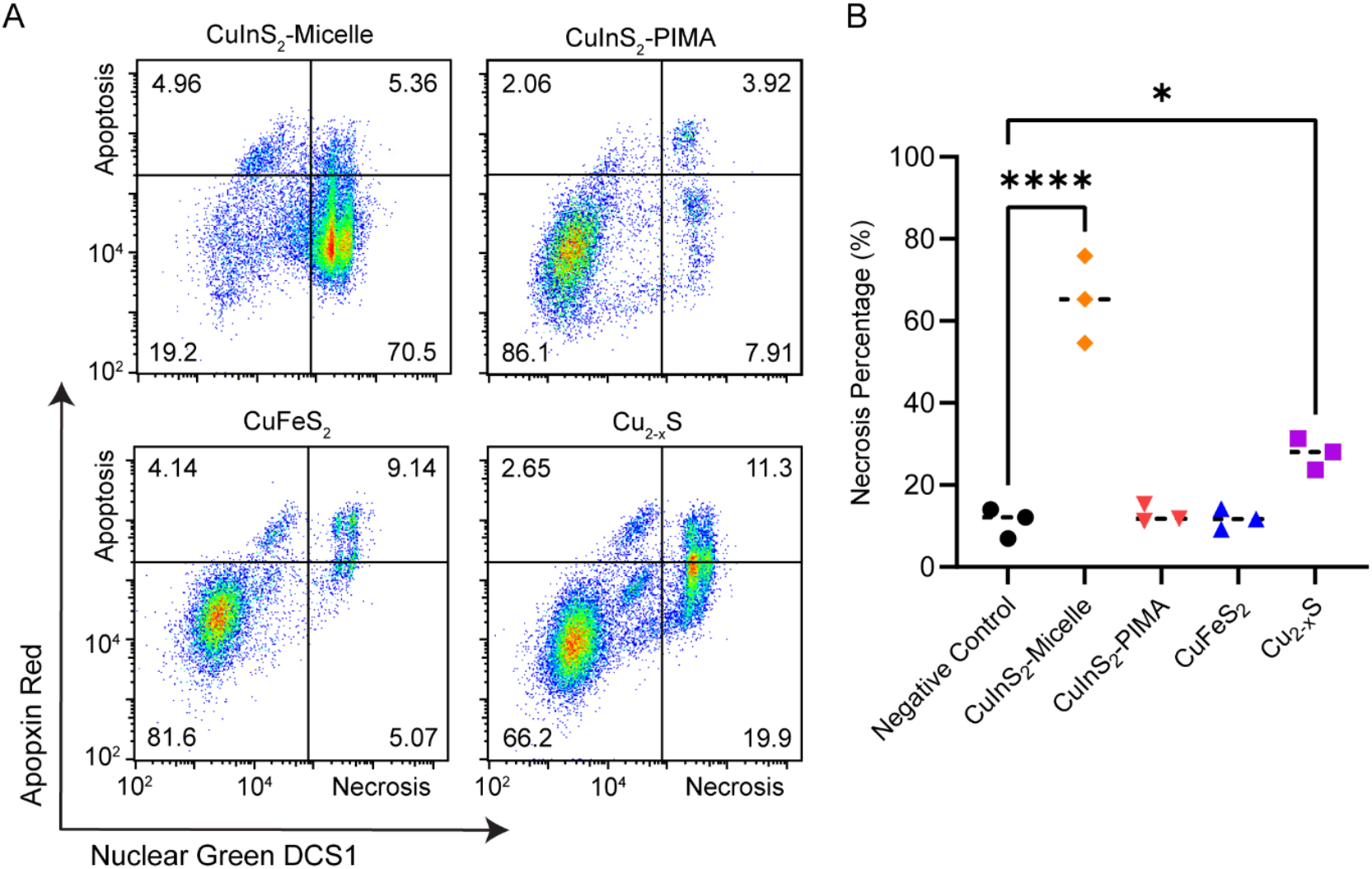
Apoptosis/necrosis assay on HepG_2_ cells after 6-hr incubation with NCs. (**A**) Flow cytometry staining of HepG_2_ cells with apoptosis and necrosis indicators Apopxin Red and Nuclear Green DSC1, respectively, following 6 hr incubation with 150 µg/mL copper chalcogenide NCs (cation concentration). (**B**). Summary and statistical analysis of the necrosis assay (n = 3). * p < 0.05, **** p < 0.0001.

**Figure 7.**
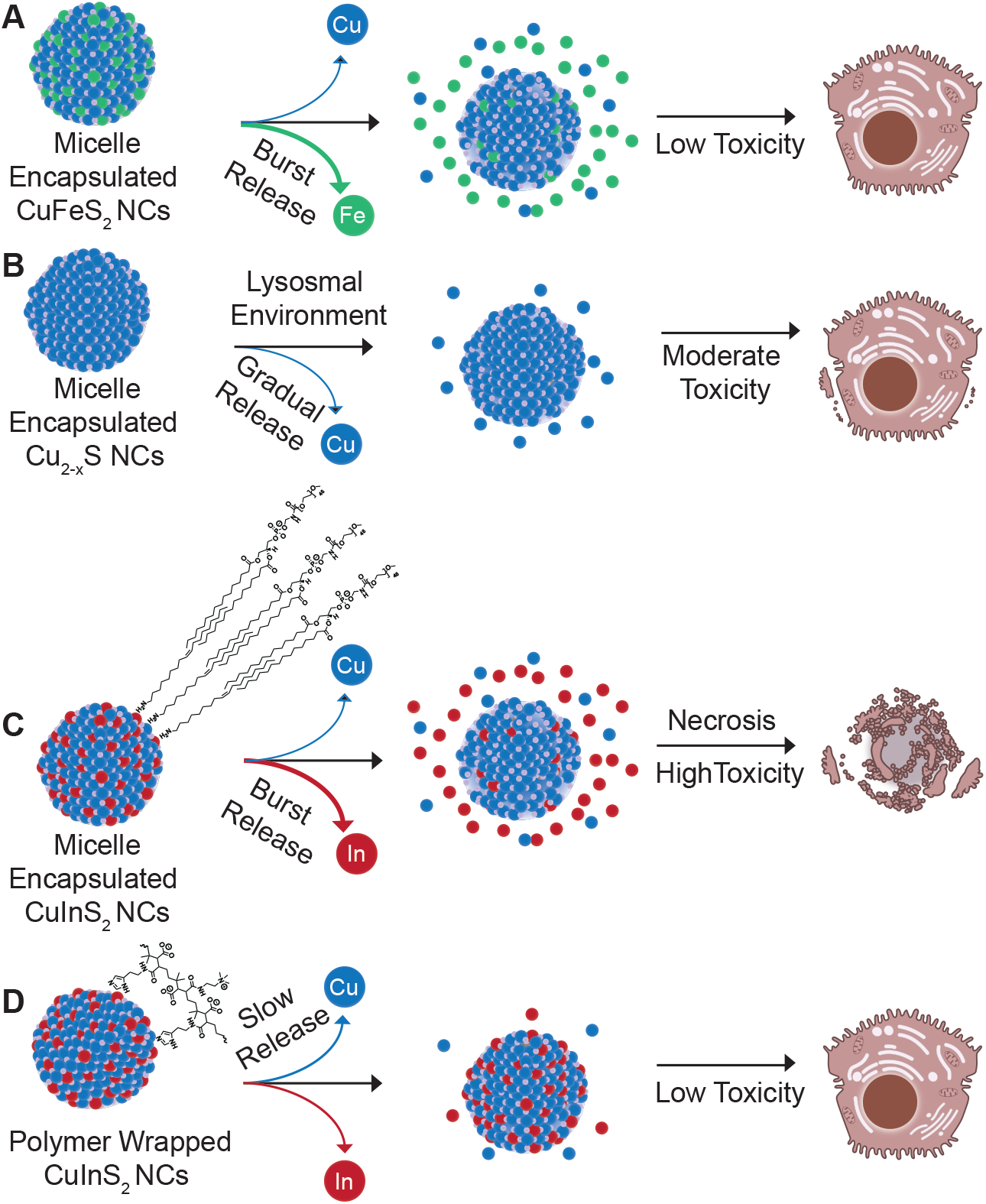
Schematic summarizing composition/coating, degradation, and toxicity results. (**A**) Burst release of iron from micelle-encapsulated CuFeS_2_ showed negligible hepatocellular toxicity, while (**B**) the release of copper from Cu_2-x_S did elicit some measurable necrosis. (**C**) CuInS_2_ NCs with the same micelle coating exhibited near immediate release of In^3+^ ions and substantial necrosis, while (**D**) wrapping the same CuInS_2_ NCs in a chelating polymer coating appears to have stabilized the particles in aqueous media mitigating hepatocellular toxicity. C and D include consolidated representations of the DSPE-PEG_2k_ and modified PIMA coatings, respectively. Schematic not to scale. Cell illustrations adapted from NIAID NIH BioArt Source (bioart.niaid.nih.gov/bioart/71 and 202)

## 3. Conclusion

In this study, we developed a quantitative approach to analyze biodegradation of small semiconductor nanocrystals and systematically compared the *in vitro* degradation and acute cytotoxicity of three compositions of micelle-encapsulated copper chalcogenide nanocrystals: CuInS_2_, Cu_2-x_S and CuFeS_2_. While the release of copper ions follows a similar gradual release pattern for each of the materials, indium and iron exhibit a rapid release shortly after dispersion in artificial lysosomal fluid. In comparative toxicity studies, CuInS_2_ exhibits the most significant acute cytotoxicity relative to the moderately toxic Cu_2-x_S and well-tolerated CuFeS_2_. This comparison identifies indium as the primary contributor to CuInS_2_ acute toxicity, primarily via necrosis, though other factors such as differences in surface reactivity between compositions may also play a role. Combining these results, we hypothesize that upon cellular uptake, CuInS_2_ undergoes lysosomal degradation, leading to a burst release of indium. The rapid release of indium, an element that is not managed by an endogenous homeostasis pathway, and the resulting high local concentration of In^3+^ likely causes cell death via necrosis. To mitigate the toxicity of CuInS_2_, we explored an alternative coating strategy using a surface chelating polymer, which appeared to slow degradation of the CuInS_2_ NCs in simulated body fluid and significantly reduced necrosis and cellular toxicity. This result highlights that in addition to the typical approach in the quantum dot community of mitigating semiconductor toxicity by encasing NCs in a stable ZnS shell, acute cytotoxicity can be reduced through careful organic coating selection with the goal of moderating the ion release rate to tolerable local concentration levels. Future work will focus on identifying organic coatings that yield favorable degradation kinetics while also improving colloidal stability over the histidine-PIMA coating used here, followed by *in vivo* validation of pharmacokinetics, biodistribution, and organ-level toxicity.

This study focused on acute cytotoxicity (6-24 hr exposures) in a hepatocellular model to understand the immediate cellular response to burst ion release from degradable nanocrystals. We selected HepG2 cells as a representative hepatic model given that the liver is the primary site of nanoparticle accumulation following systemic administration. While this focused approach enabled systematic comparison across compositions under controlled conditions, comprehensive biocompatibility assessment would require evaluation across multiple cell types (including immune cells, endothelial cells, and cells from other clearance organs), assessment of long-term toxicity and chronic exposure effects, and ultimately *in vivo* validation. Additionally, factors beyond composition and degradation, including optical excitation of these photoactive materials (which may generate reactive oxygen species), colloidal stability effects on cellular uptake, and surface properties influencing protein corona formation, may also influence nanocrystal toxicity in different contexts and warrant systematic investigation.

Overall, this study underscores the importance of thoroughly assessing composition, biodegradation, and surface coatings in the design of nanocrystals for biomedical applications. The methodological approach and insights gained from this work extend beyond copper chalcogenide nanocrystals to other metal-containing nanomaterials. For instance, degradable nanomaterials containing other non-essential metals might benefit from similar surface modification strategies to control degradation kinetics. Our quantitative degradation assay provides a valuable tool for assessing ion release from various nanoparticle compositions, especially those with smaller sizes or less dense compositions. It could be used to inform predictive toxicity models based on elemental composition and degradation profiles. Tailored degradation could be used to manage the local dose of unfavorable degradation products and improve the biocompatibility of inorganic nanoparticles, thereby expanding the range of nanoparticle compositions available for *in vivo* use.

## 4. Methods

### 4.1 Materials

Copper(II) acetylacetonate (Cu(acac)_2_, 99.9% trace metal grade), iron(III) acetylacetonate (Fe(acac)_3_, 97%), copper(I) iodide (CuI, 99.5%), indium chloride (InCl_3_, 98%), sulfur (S, 99%, trace metal grade), hexamethyldisilathiane (TMS_2_S, synthesis grade), oleylamine (OLAm, technical grade 70%), oleic acid (OA, technical grade, 90%), 1-dodecanethiol (98%), 1-octadecene (ODE, technical grade, 90%), trioctylphosphine (TOP, 97%), ethyl acetate (anhydrous), and hexane (anhydrous, 95%) were purchased from SigmaAldrich (Massachusetts, USA). Ethyl alcohol (anhydrous, >95%), chloroform (HPLC grade), dimethyl sulfoxide (DMSO, anhydrous, 99%), high purity copper standard (1,000 ppm, in 5% HNO_3_), high purity iron standard (1,000 ppm, in 5% HNO_3_), high purity indium standard (1,000 ppm, in 5% HNO_3_) and nitric acid (trace metal grade) were purchased from Fisher Scientific (New Hampshire, USA). 1,2-distearoyl-sn-glycero-3-phosphoethanolamine-N-[methoxy(polyethylene glycol)2000] (DSPE-PEG2k) was purchased from Avanti Polar Lipids (Alabama, USA).

### 4.2 Synthesis of copper chalcogenide nanocrystals

Copper indium sulfide (CuInS_2_) nanocrystals were synthesized according to previously reported methods with slight modifications.^26,50^ Specifically, in an argon-filled glovebox, 0.5 mmol (95 mg) CuI and 0.5 mmol (110 mg) InCl_3_ were added to a 100 mL round-bottomed flask along with 2.5 mL TOP, 5 mL ODE, and 3 mL OLAm. The flask was connected to a Schlenk line and heated to 95°C under vacuum until dissolved. Subsequently, the flask was backfilled with argon and heated to 170°C. Once the cation solution in the flask reached 170°C, a syringe containing 2.5 mL of 0.2 M TMS_2_S in ODE (previously prepared in the glovebox) was bolus injected. The reaction proceeded at 150°C for 20 min before heat was removed. The particle solution was degassed and transferred to the argon-filled glovebox for long-term storage.

Chalcopyrite copper iron sulfide (CuFeS_2_) nanocrystals were synthesized similarly to previously published protocols.^11,12^ In a glovebox, 0.5 mmol (130 mg) Cu(acac)_2_, 0.5 mmol (175 mg) Fe(acac)_3_, and 6.65 mL OA were added to a 100 mL round-bottomed flask. On a Schlenk line, the mixture was heated to 120°C under vacuum for 30 min until dissolved. The flask was backfilled with argon and heated to 180 °C. In a separate flask, 0.2 M sulfur in oleylamine (S/OLAm) was dissolved at 80°C under argon for 30 min. Once the cation flask reached 180 °C, 2 mL of 1-dodecanethiol was rapidly injected into the flask followed by 5 mL of S/OLAm injected dropwise over ∼30 s. The reaction was allowed to proceed for 3 min at 180°C before cooling to room temperature. The resulting solution was transferred to the argon-filled glovebox for long-term storage.

Copper sulfide (Cu_2-x_S) nanocrystals were synthesized with a method modified from literature.^51^ Specifically, 0.4 mmol (105 mg) Cu(acac)_2_ were dissolved in 7 mL oleic acid and heated to 240°C. Subsequently, 1 mL 0.2 M S/OLAm was bolus injected into the mixture followed by a second bolus injection of 3 mL of OLAm. The reaction was kept at temperature for 60 s before heat was removed, stopping the particle growth. After synthesis, nanoparticles were stored protected from light in an argon-filled glovebox.

### 4.3 Nanocrystal Characterization

Following synthesis, NCs were characterized by absorbance spectroscopy, X-ray diffraction (XRD) spectroscopy, and transmission electron microscopy (TEM). For absorbance measurements, the particles were cleaned via precipitation with a 1:3 hexane:ethanol mixture; the aggregate was pelleted via centrifugation and resuspended in hexane. Absorbance spectra were recorded with a Jasco V-770 UV-Vis-NIR spectrometer.

NC structures were examined using a D2 Phaser XRD analyzer (Bruker, MA, USA) with a coupled θ2-θ scan. Particles suspended in hexane were dropcast onto a zero-background silicon sample holder for the measurements, and XRD data were analyzed with an open-source software Fityk using reference lines obtained from the International Centre for Diffraction Data (ICDD) database.

To prepare the samples for TEM, nanoparticles were washed with hexane/ethanol mixture multiple times to remove excess ligands before resuspension in hexane. Nanocrystals suspended in hexane were dropcast onto ultrathin carbon film-coated copper TEM grids. After the hexane evaporated, grids were washed with drops of acetone and ethanol before overnight storage in a dry box. A Tecnai Osiris TEM (FEI, USA) was used for the measurements. TEM and high-resolution transmission electron microscopy (HR-TEM) images were recorded with a 300 kV electron beam.

### 4.4 Surface coatings

Chalcogenide nanocrystals were micelle encapsulated for aqueous transfer using DSPE-PEG_2k_.^33^ Briefly, particles were suspended in ∼10 mL chloroform together with a 4.5-fold mass of DSPE-PEG. The mixture was evaporated with a rotary evaporator at 65°C. Ultrapure water pre-heated to the same temperature was added to the flask together with two clean glass marbles. After vigorous swirling, the solution was passed through a 0.22 µm syringe filter to remove aggregates. The encapsulated particles were centrifuged at 68,000 rcf for at least 8 hr in a sucrose gradient column to remove empty micelles and aggregates. The density gradient column was prepared in the ultracentrifuge tube by adding 5 mL layers of sucrose solutions ranging from 60% to 20% w/w concentration in 10% increments. Each layer of sucrose solution was added once the previous layer was frozen in -80 ºC freezer for at least 15 min. Following centrifugation, particles were extracted from the sucrose layer and buffer exchanged into pH 7.4 phosphate-buffered saline using 50 kDa centrifugal filter units and stored in the fridge.

For additional surface coating comparison, CuInS_2_ nanocrystals were coated with poly(isobutylene-alt-maleic anhydride) (PIMA) functionalized with histamine as binding group to the CuInS_2_ surface.^41^ Briefly, 10 mg functionalized PIMA was dissolved in anhydrous DMSO with brief sonication. 250 µL CuInS_2_ particles were washed with 1:3 ratio of hexane: ethanol and dissolved in 9 mL of chloroform. PIMA/DMSO was added dropwise to the CuInS_2_/chloroform solution, and the mixture was stirred overnight. Coated particles were washed with ethyl acetate before resuspension in 1 mL 100 mM sodium hydroxide and 3 mL borate buffer (pH 10.5); NCs were stored at 4°C until use.

### 4.5 Degradation assay

To analyze the biodegradation of the particles in a bio-similar environment, simulated body fluid (SBF) and artificial lysosomal fluid (ALF) were made according to previous reports.^26,29,30^ For SBF, 0.8 g of sodium chloride, 0.6 g of tris(hydroxymethyl) aminomethane (Tris), 31 mg of magnesium chloride hexahydrate, 29 mg of calcium chloride, 23 mg of potassium phosphate dibasic trihydrate, 23 mg of potassium chloride, and 7.2 mg of sodium sulfate were dissolved in 100 mL of ultrapure water and the pH titrated to 7.4 with 1 M HCl. For ALF, 2.08 g of citric acid, 0.6 g of sodium hydroxide, 0.32 g of sodium chloride, 18 mg of sodium phosphate monobasic heptahydrate, 10.6 mg of magnesium chloride hexahydrate, 3.9 mg of sodium sulfate, 5.9 mg of glycerin, 9 mg of sodium tartrate dihydrate, 8.5 mg of sodium lactate, and 8.6 mg of sodium pyruvate were dissolved in 100 mL of ultrapure water and the pH adjusted to pH 4.5. The molar concentrations of all components of each buffer is listed in Table S1. To each buffers, 0.04% sodium azide was added as a preservative for long-term storage.

For absorbance-based assessment of the chalcogenide nanocrystal stability and persistence, particles were continuously incubated in SBF or ALF at 37°C. At each timepoint, the absorbance of each NC solution was recorded with a NanoDrop spectrometer (Thermo Fisher, US). The same particle solution was then returned to the incubator and remeasured at the next timepoint. The absorbance spectra were normalized to the maximum value of that solution’s original absorbance measurement (t = 0).

For the quantitative degradation assay, 1 ml ethanol was added to 500 µL micelle-encapsulated NCs in ALF or SBF at different timepoints. Precipitated NCs were collected via centrifugation at 14,000 rcf for 10 min. Both the supernatant and pellet were collected, dried, digested with nitric acid, and diluted to 5% nitric acid concentration with ultrapure water. The ion concentration in each solution was measured with microwave plasma atomic emission spectroscopy (MP-AES) using elemental standards to generate calibration curves for copper, iron, and indium. Degradation experiments were repeated at least three times. The elemental concentrations recovered in the pellet/supernatant are shown as a percentage of the original particle concentration. The earliest timepoint (t ≈ 10 min) represents the minimum practical time required for sample preparation (ethanol precipitation and centrifugation) and analysis following the start of incubation with ALF or SBF.

### 4.6 Cell culture and assays

All cell experiments were performed using the HepG_2_ cell line purchased from American Type Culture Collection (ATCC) cultured with Eagle’s Minimum Essential Medium (Corning, Catalog No. MT10009CV) with 10% Fetal Bovine Serum (Corning, Catalog No. MT35015CV) according to the ATCC guide.

For cell viability measurements, cells were plated in 96-well plates with a cell density of 40,000 cells/well and allowed to adhere overnight. Concentrated particles in PBS were added to media at varying concentrations for a 24-hr cell incubation with n = 4 replicates. Media with the same amount of phosphate-buffered saline (PBS, pH 7.4) and no particles were used as negative controls for cell viability percentage calculations. The amount of PBS in media does not exceed 15% in all studies. A CellTiter-Glo cell viability assay (Promega, WI) was used to measure cell viability. After nanoparticle incubation, cells were washed with PBS and treated with the cell viability assay mixture following the assay protocol. After a 10 min incubation, bioluminescence was measured in a plate reader (Paradigm, Molecular Devices, CA) using a 250 ms exposure time.

Necrosis/apoptosis was assayed using a three-color fluorescence kit from Abcam (Catalog No. ab176749). Cells were plated in 12-well plate with a density of 500,000 cells per well overnight. Particles at 150 µg/mL cation concentration (Cu+In, Cu, or Cu+Fe) were added to the cells for 24-hr incubation. For apoptosis positive controls, 1 – 5 μM staurosporine was used for a 6 hr incubation. For necrosis positive controls, 90% ethanol was incubated with cells for 60 s at 37°C and immediately quenched with a 5-fold excess of phosphate-buffered saline. After incubation, media was collected from each well and adhered cells were detached using TrypLE enzyme (Gibco, Catalog No. 12604021) and quenched with the culture media. The two solutions containing floating and adherent cells, respectively, were combined and centrifuged at 500 rcf for 5 min. The supernatant containing the particles was discarded and the cells were treated with the assay kit for 30 min in the dark at room temperature. The cells were measured using a multi-channel CytoFLEX S flow cytometry analyzer (Beckman Coulter, CA, USA). Data were analyzed and plotted using CytExpert (Beckman Coulter, CA, USA).

## Supporting Information

Absorbance spectrum of 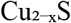 nanocrystals extending into the near-infrared; dynamic light scattering and zeta potential measurements for all nanocrystal compositions; TEM size distribution histograms; quantitative comparison of copper release from all three nanocrystal compositions in SBF and ALF; indium and iron release comparison in SBF; flow cytometry control experiments for apoptosis/necrosis assay including negative control, necrosis positive control, and apoptosis positive controls; statistical comparison of apoptosis populations across experimental groups.

## Acknowledgement

The research reported in this publication was supported by the National Institutes of Health through the National Institute of General Medical Sciences (NIH NIGMS; Award No. R01GM129437) and the National Institute of Biomedical Imaging and Bioengineering (NIH NIBIB; Award No. R21EB032647) as well as the National Science Foundation (NSF; Award No. 2403855). The content is solely the responsibility of the authors and does not necessarily represent the official views of the National Institutes of Health. We thank the Institute for Chemical Imaging of Living Systems (RRID:SCR_022681) at Northeastern University for consultation and flow cytometry support. We appreciate the Boston University Materials Science and Engineering core facility and Chemistry Department Chemical Instrumentation Center for access to their XRD and MP-AES instrumentation, respectively.

## Author Contribution

X.Z. and A.M.D. contributed to conceptualization, experimentation, data analysis and manuscript writing. G.P.K contributed to degradation and cell viability experiments. S.N. contributed to cellular assays.

## Table of Contents graphic

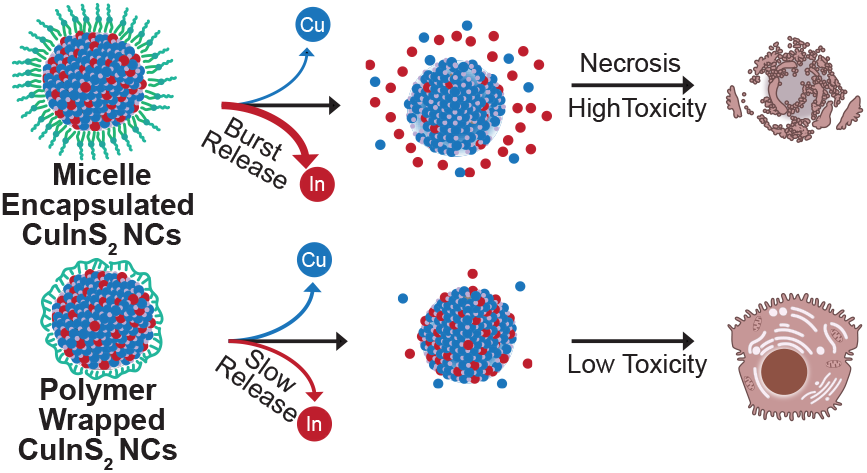

