## Supplemental Information for "Elemental Composition and Degradation Rate Impact the Biocompatibility of Copper Chalcogenide Nanocrystals"

Figure S1. Size distribution of copper chalcogenide nanocrystals

Figure S2. Absorbance spectrum of micelle-encapsulated $Cu_{2-x}S$

Figure S3. Dynamic light scattering (DLS) hydrodynamic diameter and ζ-potential measurements

Figure S4. Cation release comparison between compositions

Figure S5. Control experiment results of apoptosis/necrosis assay measured with flow cytometry

Figure S6. Summary of apoptosis results

Table S1. Ionic composition of simulated body fluid (SBF) and artificial lysosomal fluid (ALF)

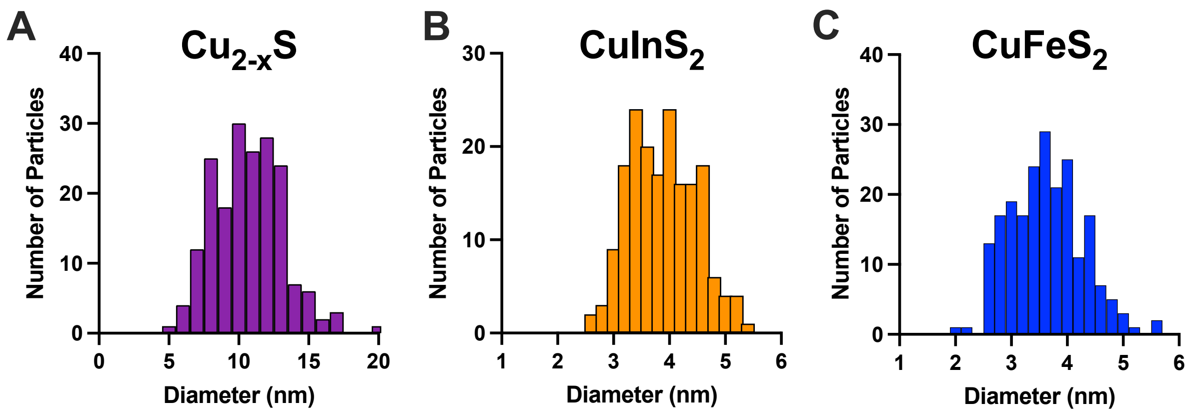


**Figure S1. Size distribution of copper chalcogenide nanocrystals** measured from TEM images. Histograms show particle size distributions for (**A**) $Cu_{2-x}S$, (**B**) $CuInS_2$, and (**C**) $CuFeS_2$. Mean ± standard deviation of particle diameters: $Cu_{2-x}S$ = 10.7 ± 2.5 nm (n = 187); $CuInS_2$ = 3.9 ± 0.6 nm (n = 182); and $CuFeS_2$ = 3.6 ± 0.7 nm (n = 213).

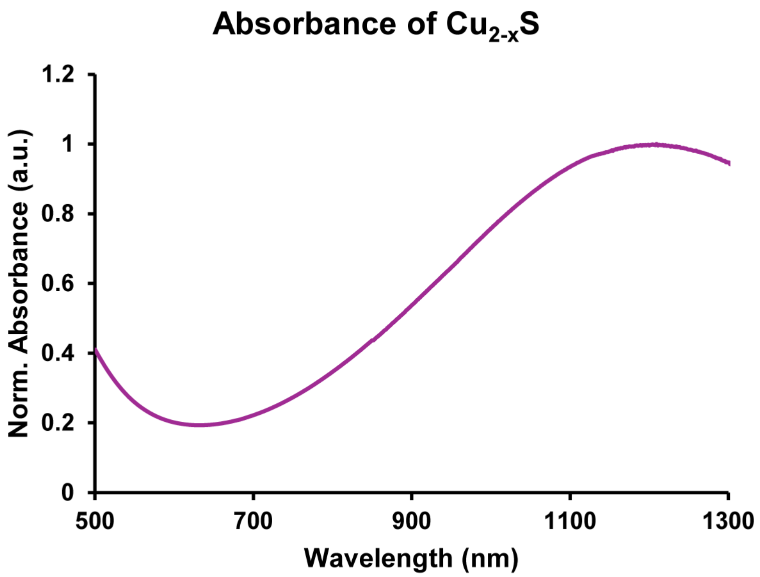


**Figure S2. Absorbance spectrum of micelle-encapsulated $Cu_{2-x}S$ nanocrystals** into the near-infrared range. Data is normalized to absorbance peak. Absorbance at wavelengths > 1300 nm is not shown due to large artifacts from water absorbance.

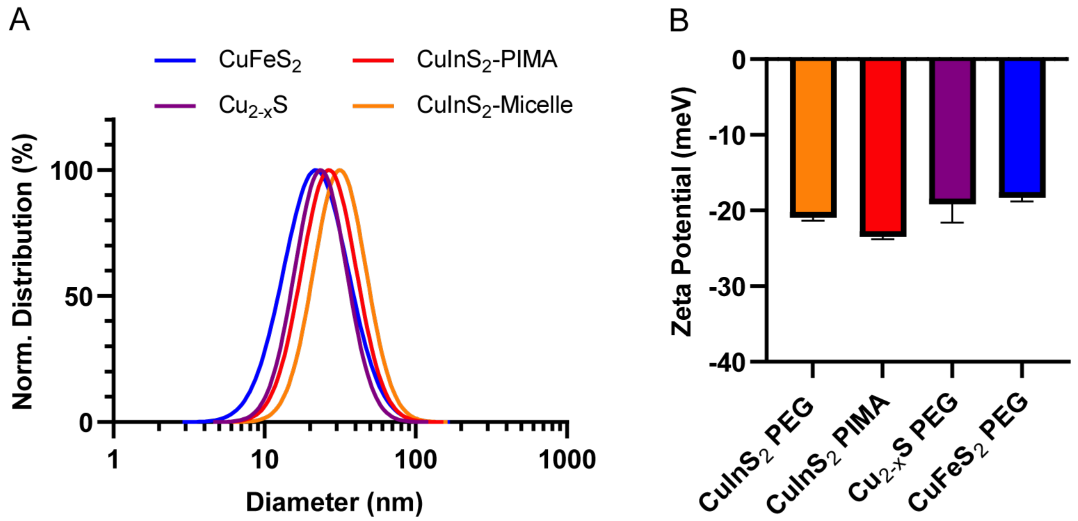


**Figure S3. Dynamic light scattering (DLS) hydrodynamic diameter and ζ-potential measurements.** (**A**) Number-weighted DLS measurements on log-normal plot. Average hydrodynamic diameter based on number-weighted peaks: $CuFeS_2$ (22.8± 1.8 nm); $Cu_{2-x}S$ (26.0 ± 4.9 nm); $CuInS_2$-Micelle (36.0 ± 4.4 nm); $CuInS_2$-PIMA (30.8 ± 1.2 nm). (**B**) ζ-potential measurements of particles. DLS and Zeta were measured with NanoBrook 90Plus PALS (Brookhaven, NH) with n = 3 repeats.

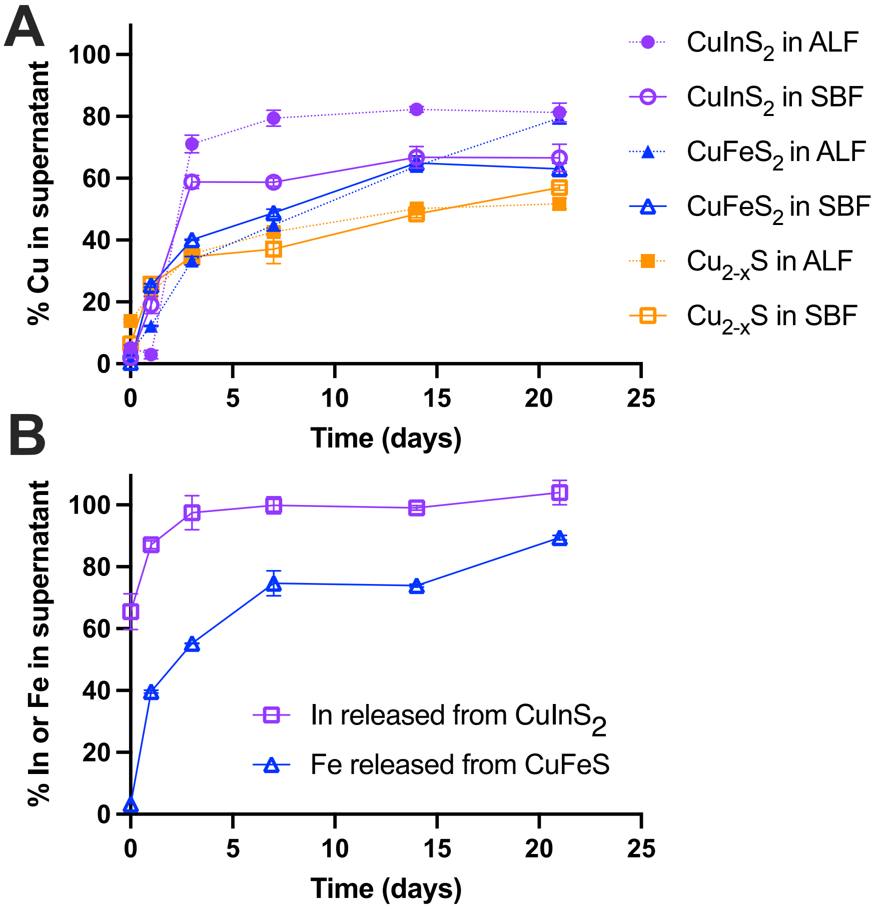


**Figure S4. Cation release comparison between compositions.** (**A**) Release of copper into the supernatant upon incubation of micelle-encapsulated copper chalcogenides in SBF and ALF. (**B**) Release of indium and iron into the supernatant upon incubation of micelle-encapsulated NCs in SBF.

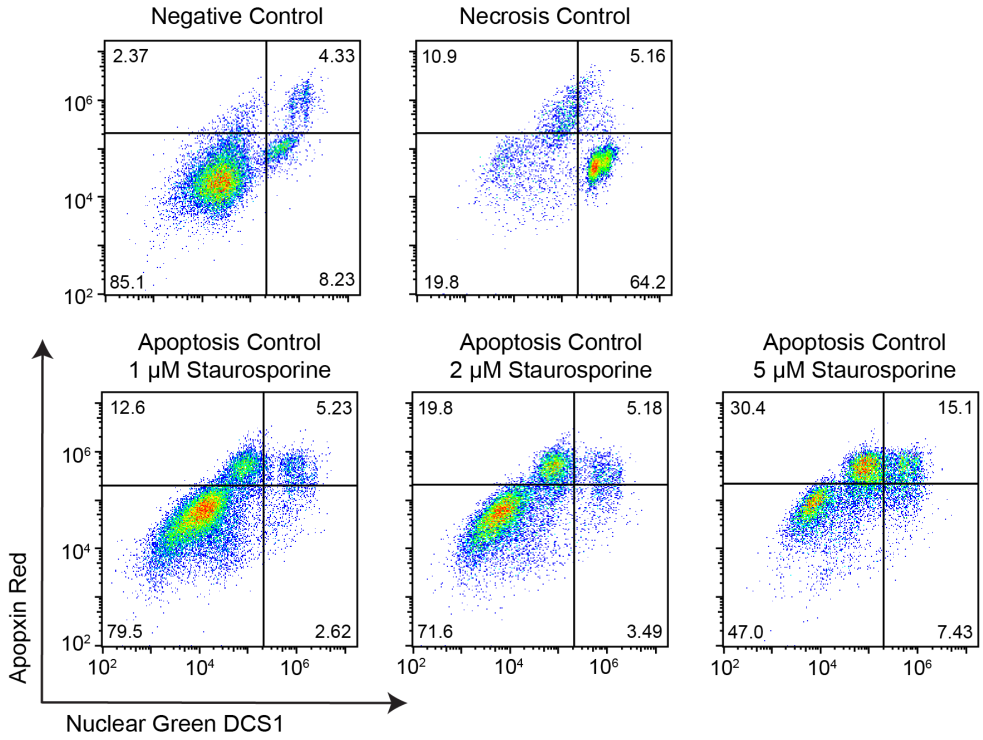


**Figure S5. Control experiment results of apoptosis/necrosis assay measured with flow cytometry.** Particles were incubated with pH 7.4 PBS buffer for negative control; 90% ethanol for 60 seconds for necrosis positive control; 1 – 5 μM staurosporine for 24 hr for apoptosis positive control.

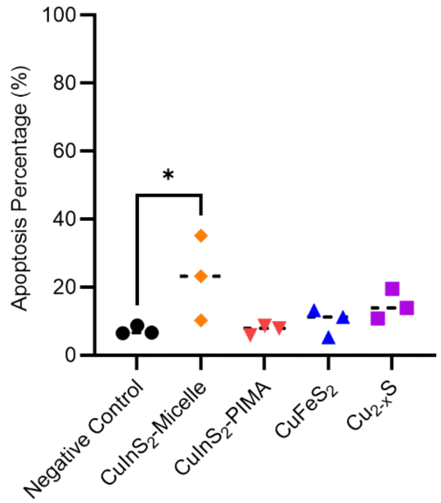


**Figure S6. Summary of apoptosis results.** Comparison between different experimental groups and negative control. *P < 0.05.

**Table S1. Ionic composition of simulated body fluid (SBF)[1,2] and artificial lysosomal fluid (ALF),[3]** adapted from previous reports**.**

| ***Ions*** | ***Simulated Body Fluid (SBF) (mM)*** | ***Artificial Lysosomal Fluid (ALF) (mM)*** |
|---|---|---|
| *$Na^+$* | 138 | 208.4 |
| *$Ca^{2+}$* | 2.6 | - |
| *$Mg^{2+}$* | 1.5 | 0.5 |
| *$K^+$* | 5 | - |
| *$Cl^-$* | 148.8 | 55.8 |
| *$SO_4^{2-}$* | 0.5 | 0.3 |
| *$HPO_4^{2-}$* | 1 | - |
| *$H_2PO_4^-$* | - | 0.7 |
| *Tris* | 50 | - |
| *Hydrochloric acid* | 40 | - |
| *Tartrate* | - | 0.4 |
| *Lactate* | - | 0.8 |
| *Pyruvate* | - | 0.8 |
| *Citric Acid* | - | 108 |
| ***pH*** | 7.4 | 4.5 |
